# TNIP1 and Autophagy Receptors regulate STING Signaling

**DOI:** 10.1101/2025.04.21.649822

**Authors:** Eric N. Bunker, Tara D. Fischer, Peng-Peng Zhu, François Le Guerroué, Richard J. Youle

**Author notes:** Co-first authors (Eric N. Bunker, Tara D. Fischer). Present Address: Single Cell Biomarkers UTechS, Institut Pasteur, Université Paris Cité, Paris, France (François Le Guerroué).

## Abstract

Activation of the cGAS-STING pathway stimulates innate immune signaling as well as LC3B lipidation and ubiquitylation at Golgi-related vesicles upon STING trafficking. Although ubiquitylation at these subcellular sites has been associated with regulating NF-κB-related innate immune signaling, the mechanisms of Golgi-localized polyubiquitin chain regulation of immune signaling is not well understood. We report here that the ubiquitin– and LC3B-binding proteins, TNIP1 and autophagy receptors p62, NBR1, NDP52, TAX1BP1, and OPTN associate with STING-induced ubiquitin and LC3B-labeled vesicles, and that p62 and NBR1 act redundantly in spatial clustering of the LC3B-labeled vesicles in the perinuclear region. We also find that while TBK1 kinase activity is not required for the recruitment of TNIP1 and the autophagy receptors, it also plays a role in sequestration of the LC3B-labeled vesicles. The ubiquitin binding domains, rather than the LC3B-interacting regions, of TNIP1 and OPTN are specifically important for their recruitment to Ub/LC3B-associated perinuclear vesicles, while OPTN is also recruited through a TBK1-dependent mechanism. Functionally, we find that TNIP1 and OPTN play a role in STING-mediated innate immune signaling, with TNIP1 acting as a significant negative regulator of both NF-κB– and Interferon-mediated gene expression. Together, these results highlight autophagy-independent mechanisms of autophagy receptors and TNIP1 with unanticipated roles in regulating STING-mediated innate immunity.

## Introduction

Activation of the antiviral defense protein STING initiates diverse cellular responses, including interferon signaling, NF-κB signaling, and LC3B lipidation at Golgi-related membranes (Hopfner & Hornung, 2020; B. Zhang et al., 2025). STING binding to cyclic dinucleotide 2’3’ cGMP-AMP (cGAMP) induces the oligomerization of STING dimers and their translocation from the ER to the Golgi apparatus (Ergun et al., 2019; Ishikawa et al., 2009; Saitoh et al., 2009; Shang et al., 2019). Following trafficking to the Golgi and formation of higher order STING oligomers, the kinase TBK1 binds to STING, and is activated to phosphorylate STING and the transcription factor IRF3 (Tanaka & Chen, 2012; C. Zhang et al., 2019). Phosphorylated IRF3 then initiates the transcription of Interferon β and downstream interferon-stimulated signaling. TBK1 also acts redundantly with IKKε in propagating NF-κB signaling mediated by STING (Balka et al., 2020). Independently of TBK1, LC3B lipidation induced by STING activation is mediated by the V-ATPase-ATG16L1 axis at Golgi-related vesicles (Fischer et al., 2020; Gui et al., 2019) following collapse of the proton gradient across Golgi membranes (Liu et al., 2023; Xun et al., 2024). This process is similar to the conjugation of ATG8s at single membranes (CASM) induced by chemical and pathogen-related endo-lysosomal damage (Durgan et al., 2021). We have recently demonstrated that polyubiquitin chains are co-localized with LC3B and activated STING in the perinuclear region of cells near the Golgi (Fischer et al., 2024). STING activation induces the E3 ligase HOIP to synthesize M1-linked polyubiquitin chains that stimulates NF-κB signaling (Fischer et al., 2024); however, the mechanism of the Golgi-related localization of ubiquitin impacts innate immunity is not well understood.

Ubiquitin binding proteins are key regulators of downstream ubiquitin signaling (Oh et al., 2025). Several ubiquitin binding proteins are known to also bind LC3B and function in sequestration and degradation of ubiquitylated cargo in selective autophagy (Johansen & Lamark, 2020; Kirkin & Rogov, 2019). However, many of these ubiquitin binding proteins are also implicated in regulating innate immune signaling through unclear mechanisms (Deretic, 2021; Shamilov & Aneskievich, 2018; Slowicka et al., 2016). Here, we aimed to determine if autophagy-related ubiquitin binding proteins play a role at LC3B– and Ub-associated Golgi-vesicles in the STING pathway.

## Results

We assessed the spatial localization of a panel of ubiquitin binding proteins with LC3 Interacting Region (LIR) domains – TNIP1 and p62, NBR1, NDP52, TAX1BP1, and OPTN (also known as ‘autophagy receptors’ and further referred as such) by immunofluorescent labeling, in HeLa cells stably expressing STING at low levels (HeLa^STING^) and mScarletI-LC3B after treatment with the STING agonist diABZI (Figure 1). diABZI treatment induced a re-localization of each autophagy receptor and TNIP1 to form foci in the perinuclear region of cells (Figure 1, A-G) that robustly colocalize with LC3B foci (Figure 1, A-F and H), showing that these soluble ubiquitin binding proteins are recruited to LC3B-associated vesicles upon STING activation.

**Figure 1.**
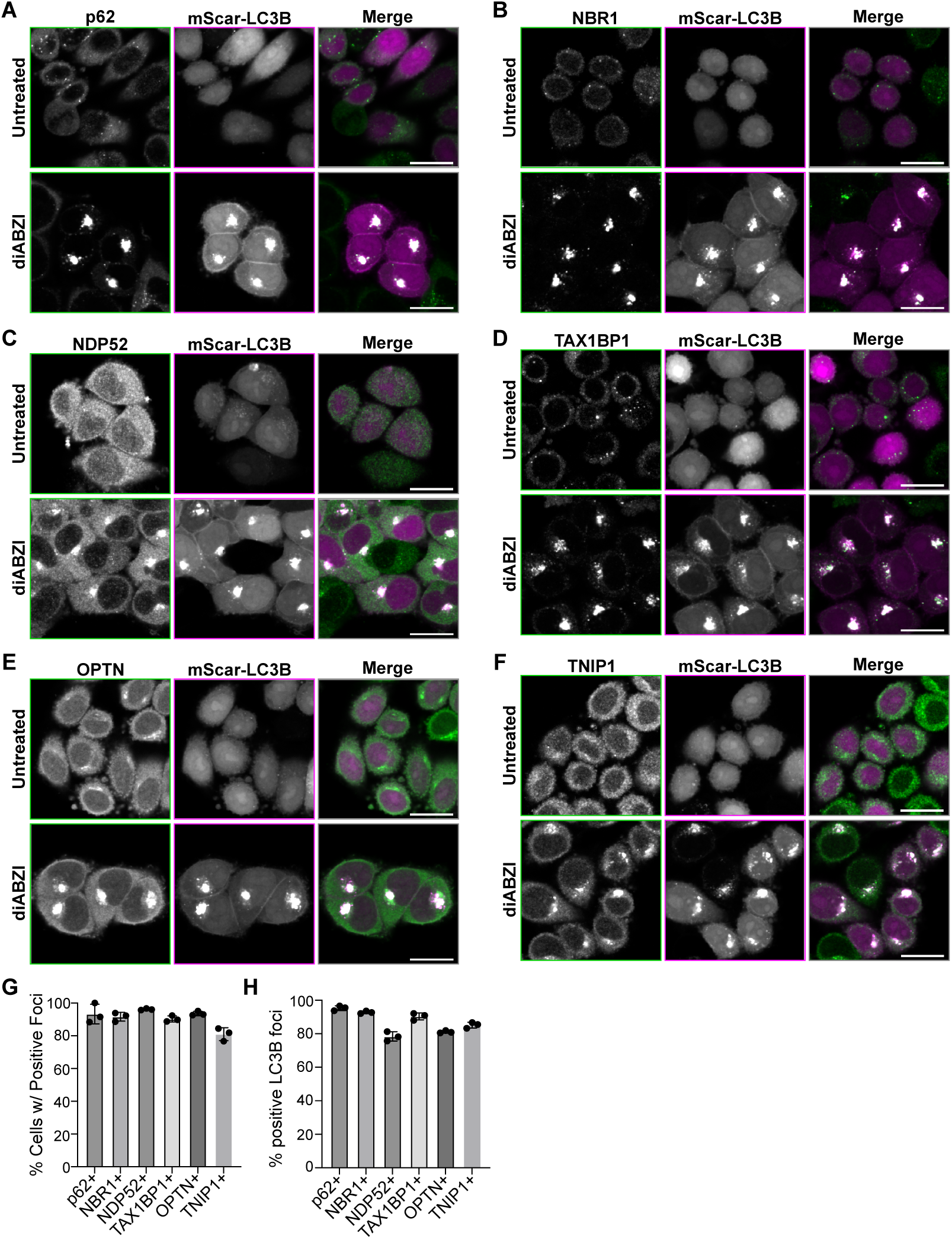
Autophagy receptors and TNIP1 are recruited to perinuclear LC3B foci upon STING activation. **A-F**) Representative spinning disk confocal images depicting cells either untreated or treated with diABZI for 6 hr before fixation. Green represents immunostaining of indicated protein and magenta represents mScarletI-LC3B. Scale bar indicates 20 µm. **G-H)** Quantification of images in panels A-E depicting the fraction of LC3B positive cells also positive for foci of each indicated protein or LC3B foci positive for indicated protein. Over 1000 cells were imaged for each condition.

Time-lapse imaging of LC3B foci formation following diABZI activation of STING shows that fluorescently tagged LC3B first forms dispersed puncta in the perinuclear region of cells at early timepoints (1 hour), that cluster into a singular bright focus or a few large foci within each cell (Figure 2A). To quantify the different spatial distributions of dispersed puncta and foci, we developed a method of analysis in which the weighted centroid of signal (LC3B or other fluorescently labeled proteins) within each cell is found, and then the relative fluorescence intensity of the signal in relation to its distance from the centroid is measured. This analysis for mEGFP-LC3B signal displays a peak followed by a sharp decline in intensity in WT cells at later timepoints following diABZI treatment (Figure 2B; bright green), consistent with foci with defined edges, whereas early timepoints display a smaller peak and broader slope from the centroid, consistent with the dispersed pattern (Figure 2B; dark green). Because autophagy receptors are important for sequestering cargo for selective autophagy (Deosaran et al., 2013; Kirkin & Rogov, 2019; Narendra et al., 2010; Sun et al., 2018; Turco et al., 2021; Zaffagnini et al., 2018), we tested whether these autophagy receptors are important for the ‘clustering’ of LC3B into foci upon STING activation. Compared to WT HeLa^STING^ cells, LC3B and ubiquitin in cells lacking p62, NBR1, NDP52, TAX1BP1, and OPTN (Lazarou et al., 2015) (5KO HeLa^STING^), formed dispersed puncta in the perinuclear region, but not foci upon diABZI treatment (Figure 2, C-E), indicating that the autophagy receptors may be involved in the clustering of LC3B and Ub positive Golgi-related vesicles following STING activation. Both p62 and NBR1 are particularly known to induce clustering of cargo in selective forms of autophagy (Sun et al., 2018; Turco et al., 2021; Zaffagnini et al., 2018) through their PB1 domains, as well as the amphipathic α-helix in NBR1 that allows sequestration of vesicles (Deosaran et al., 2013; Mardakheh et al., 2010). To determine whether p62 or NBR1 are specifically required for LC3B and Ub foci formation, we compared LC3B distribution in p62KO and NBR1KO HeLa^STING^ cells to that in WT HeLa^STING^ cells following diABZI treatment; however, no change was found with either KO (Figure S1, A-C). To assess if p62 and NBR1 may be redundant in this regard, we also compared p62/NBR1 DKO HeLa^STING^ cells to WT. In p62/NBR1 DKO HeLa^STING^ cells, we found LC3B displayed dispersed puncta following diABZI treatment (Figure 2, F-G) similarly to that of 5KO cells, which could be rescued by expression of either mEGFP-p62 or –NBR1 (Figure 2, F-G). Together, these data indicate that p62 and NBR1 are redundantly capable of clustering LC3B/Ubiquitin associated vesicles following STING activation.

**Figure 2.**
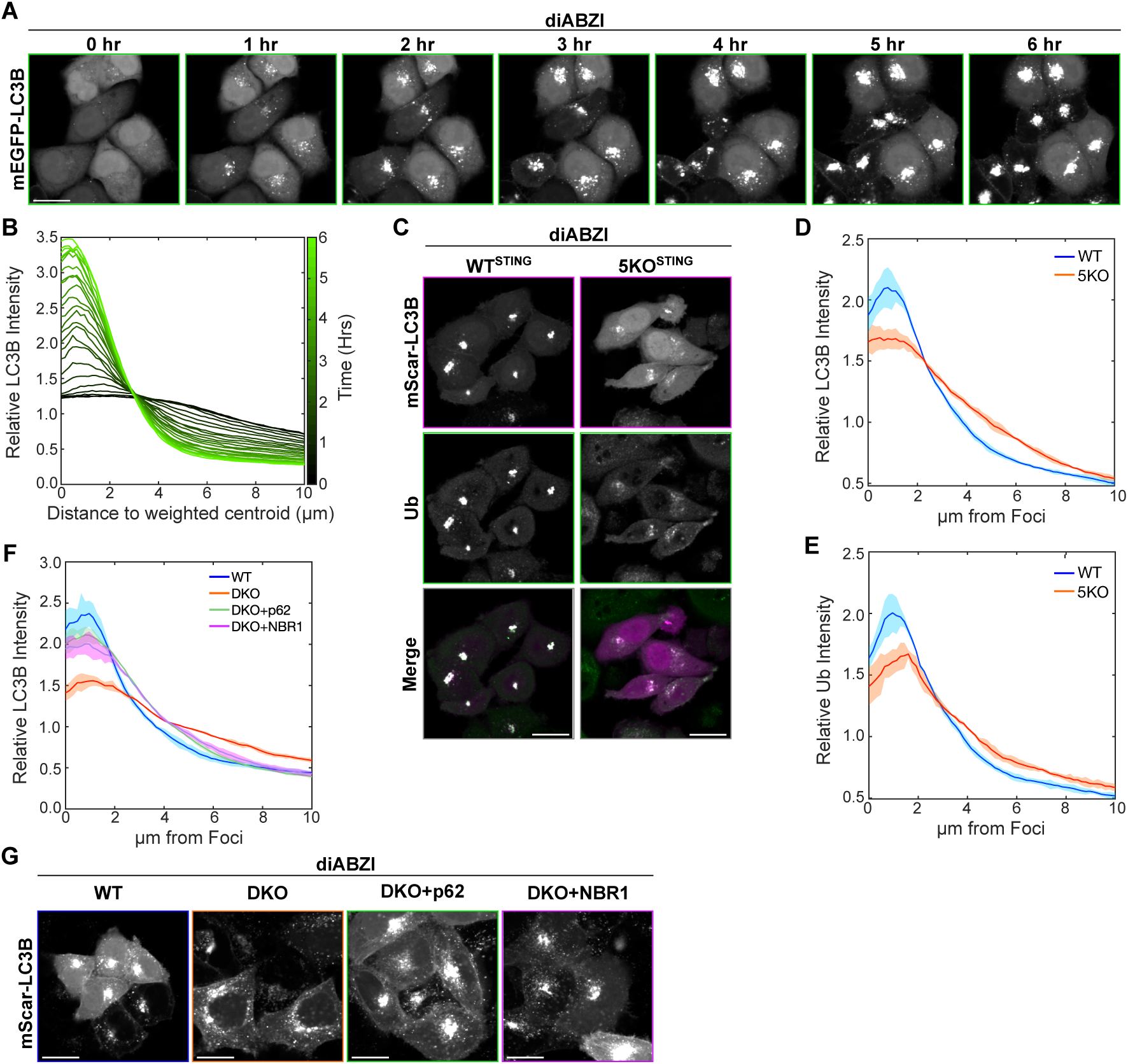
p62 and NBR1 act redundantly to cluster STING-induced LC3B and Ubiquitin into perinuclear foci. **A)** Representative spinning disk confocal images of WT HeLa^STING^ cells expressing mEGFP-LC3B treated with diABZI. Scale bar indicates 20 µm. **B)** Quantification of normalized intensity of LC3B signal within each cell plotted against distance to centroid of LC3B signal from cells displayed in panel A. Color indicates timepoint measured, where darker green is earlier after diABZI treatment and lighter green is closer to 6 hr after diABZI treatment. Over 100 cells were used for quantification. **C)** Representative images of WT or 5KO HeLa^STING^ (lacking *p62*, *NBR1*, *NDP52*, *TAX1BP1*, and *OPTN*) with mScarlet-LC3B in magenta and ubiquitin in green. Scale bar indicates 20 µm. **D-E)** Quantification of normalized intensity of LC3B or ubiquitin signal within each cell plotted against distance to centroid of LC3B signal from cells displayed in panel C. Over 1000 cells were imaged for each condition. **F)** Quantification of normalized intensity of LC3B signal within each cell plotted against distance to centroid of LC3B signal from cells displayed in panel G. Over 1000 cells were imaged for each condition. **G)** Representative images of LC3B after 6 hr diABZI treatment in WT or DKO HeLa^STING^, (lacking *p62* and *NBR1*), DKO HeLa^STING^ expressing mEGFP-p62, or DKO HeLa^STING^ expressing mEGFP-NBR1. Scale bars indicate 20 µm.

TBK1 plays a major role in STING signaling and is also known to function in selective autophagy by phosphorylating autophagy receptors (Heo et al., 2015; Matsumoto et al., 2015; Moore & Holzbaur, 2016; Pilli et al., 2012; Richter et al., 2016; Wild et al., 2011; Zhou et al., 2023). Interestingly, we discovered that LC3B formed dispersed puncta, rather than foci, also in TBK1KO cells (Figure 3, A-B) similarly to early time points of diABZI treatment (Figure 2, A-B), 5KO cells (Figure 2, C-D), and p62/NBR1 DKO cells (Figure 2, F-G). Although previous reports have demonstrated that TBK1 is not required for LC3B lipidation (Gui et al., 2019), these results suggest that TBK1 is involved in the clustering of LC3B-associated membranes following STING activation. To determine whether TBK1 kinase activity is required for LC3B clustering into foci, we treated cells with the TBK1 kinase inhibitor, GSK8612 (GSK), as well as diABZI, and found a substantial decrease in LC3B clustering following STING activation (Figure 3, C-F). The use of a TBK1 inhibitor also allowed us to assess the dynamics of formation and stability of LC3B foci. We found that the dispersed LC3B puncta clustered into foci following removal of the TBK1 inhibitor after 6 hours of diABZI + GSK treatment (Figure 3, C-F), while LC3B foci were retained if the TBK1 inhibitor was added at 6 hours post-diABZI treatment (Figure 3, C-F). Together these data demonstrate that TBK1 kinase activity is involved in the clustering of LC3B-lipidated vesicles, but once LC3B foci have formed, TBK1 activity is dispensable.

**Figure 3.**
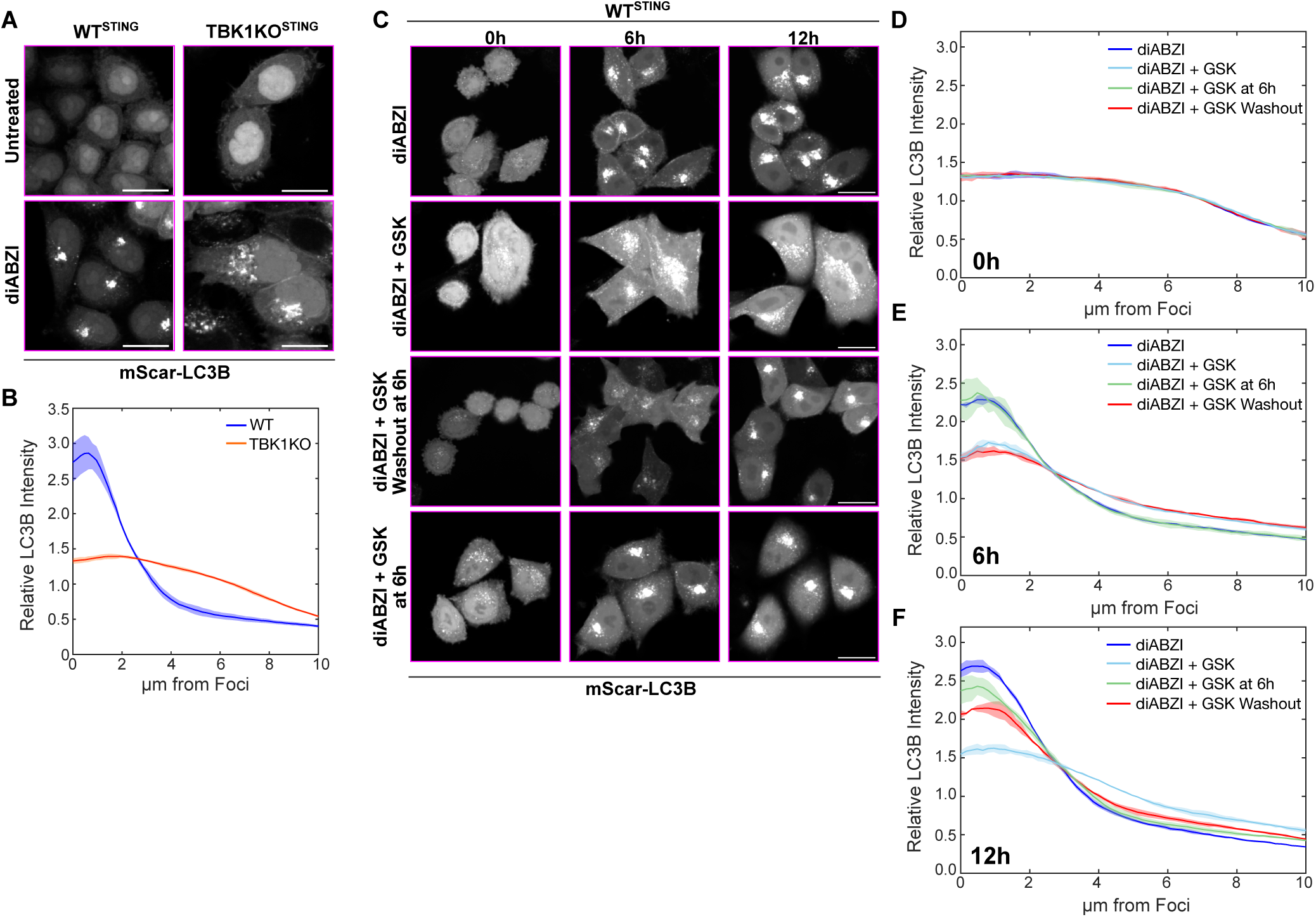
TBK1 kinase activity promotes LC3B clustering into perinuclear foci. **A)** Representative spinning disk confocal images of mScarlet-LC3B either untreated or after 6 hr diABZI treatment in WT and TBK1KO HeLa^STING^ cells. Scale bars indicate 20 µm. **B)** Quantification of normalized intensity of LC3B signal within each cell plotted against distance to centroid of LC3B signal from cells displayed in panel A. Over 1000 cells were imaged for each condition. **C)** Representative spinning disk confocal images of mScarletI-LC3B either immediately after (left column), 6 hr after (middle column), or 12 hr after (right column) diABZI treatment in WT and TBK1KO HeLa^STING^ cells. TBK1 kinase inhibitor GSK added for all 12 hr (second row), first 6 hr then washed out (third row), or added for the last 6 hr (bottom row). Scale bars indicate 20 µm. **D-F)** Quantification of normalized intensity of LC3B signal within each cell plotted against distance to centroid of LC3B signal from cells displayed in panel C, measured at 0 hr (D), 6 hr (E) or 12hr (F). Over 1000 cells were imaged for each condition.

Autophagy receptor recruitment and activity can be regulated by TBK1 (Heo et al., 2015; Lazarou et al., 2015; Moore & Holzbaur, 2016; Richter et al., 2016; Vargas et al., 2019). As TBK1 also interacts with activated STING, we hypothesized that TBK1 and its kinase activity may be required for autophagy receptor and TNIP1 recruitment to LC3B foci. Combined treatment of WT HeLa^STING^ cells with diABZI and the TBK1 kinase inhibitor GSK revealed no defect in recruitment of any of the endogenous autophagy receptors or TNIP1 to LC3B structures upon TBK1 kinase inhibition (Figure S2, A-G). OPTN directly binds to TBK1 regardless of kinase activity (Li et al., 2016; Morton et al., 2008; Wild et al., 2011), so we also compared recruitment of each autophagy receptor in TBK1KO relative to WT HeLa^STING^ cells. There was no change in recruitment of p62, NBR1, NDP52, TAX1BP1, and TNIP1 in TBK1KO cells but a substantial decrease in OPTN recruitment in TBK1KO cells following diABZI treatment was detected (Figure 4, A-G). These results indicate that the interaction between TBK1 and OPTN may be important for OPTN recruitment to LC3B foci following STING activation, independent of TBK1 kinase activity. As STING directly binds TBK1 upon activation (C. Zhang et al., 2019), we sought to determine if the interaction between STING and TBK1 is important for recruitment of OPTN to LC3B-labeled vesicles. We mutated the TBK1 interacting region of STING (L374A) and compared OPTN recruitment to LC3B following diABZI treatment in cells stably overexpressing WT STING or STING-L374A (C. Zhang et al., 2019; Zhao et al., 2019). Similar to the results in TBK1KO cells, there was a substantial reduction in OPTN recruitment to LC3B foci in cells expressing STING-L374A (Figures 4H and S2H), implicating the STING-TBK1 interaction for recruiting OPTN in this pathway.

**Figure 4.**
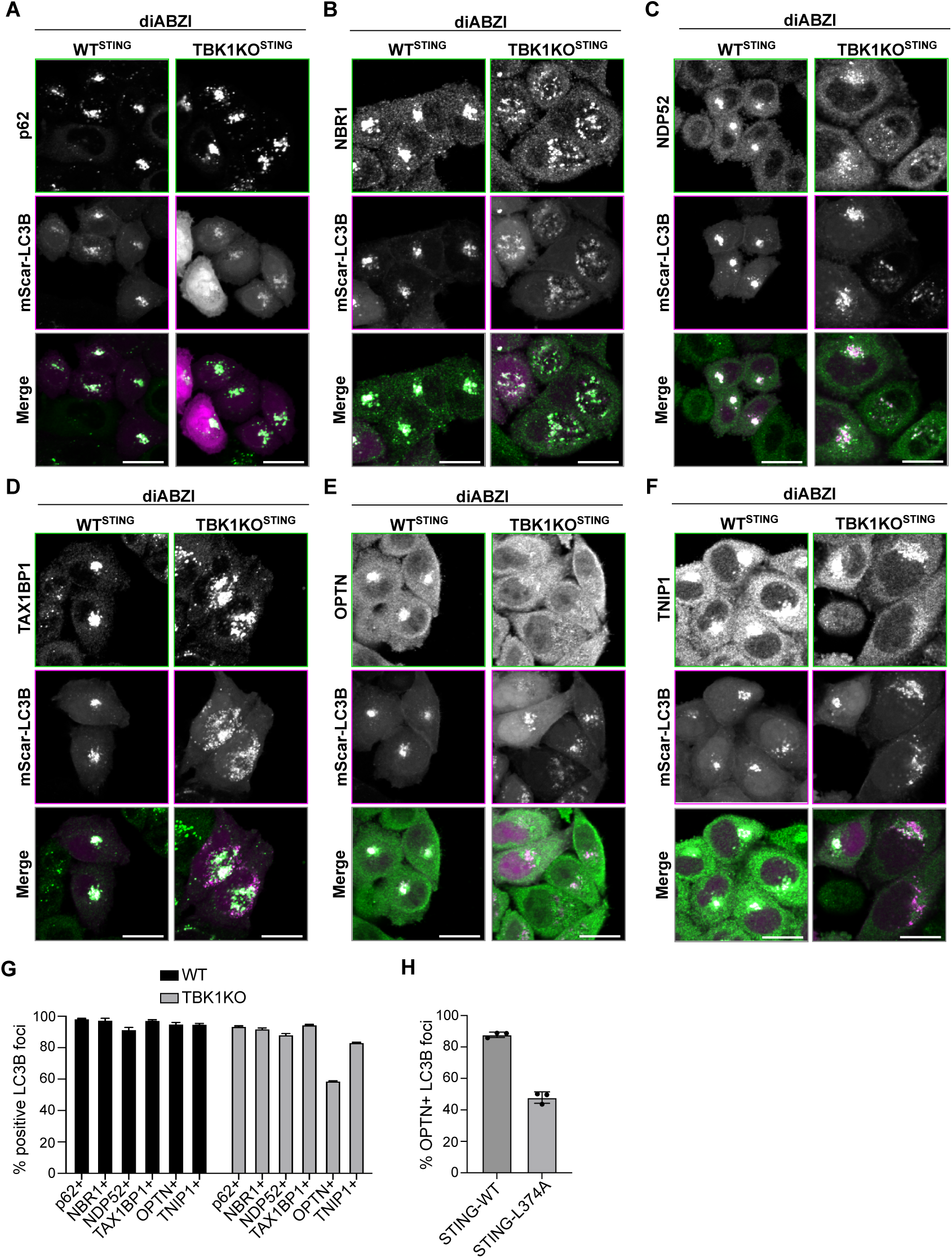
TBK1 is important for STING-induced OPTN recruitment to LC3B foci but is dispensable for recruitment of other autophagy receptors and TNIP1. **A-F)** Representative spinning disk confocal images of either WT or TBK1KO HeLa^STING^ cells treated with diABZI for 6 hr before fixation. Green represents immunostaining of indicated protein and magenta represents mScarletI-LC3B. Scale bar indicates 20 µm. **G)** Quantification of images in panels A-E depicting fraction of LC3B foci positive for indicated proteins. Over 1000 cells were imaged for each condition. **H)** Quantification of images in Figure S2G, depicting fraction of LC3B foci positive for OPTN in cells expressing WT STING or STING_L374A_. Over 1000 cells were imaged for each condition.

STING activation involves STING trafficking and oligomerization at post-Golgi vesicles where it may form an ion channel (Liu et al., 2023; Xun et al., 2024), inducing disruption of pH gradients and LC3B lipidation (Fischer et al., 2020) similarly to that of membrane damaging agents linked to CASM (Figueras-Novoa et al., 2024). To determine if autophagy receptors and TNIP1 are recruited to LC3B-associated single membrane vesicles upon membrane damage of different organelles by CASM-related stimuli, we treated cells with the ionophore Monensin (Florey et al., 2015), a membranolytic polymer that damages lysosomes, LLOMe (Cross et al., 2023; Thiele & Lipsky, 1990), and a small molecule reported to damage the Golgi, AMDE-1 (Y. Gao et al., 2016). Upon Monensin treatment, p62, NBR1, TAX1BP1 and NDP52 are robustly recruited to the LC3B puncta/foci formed, whereas OPTN and TNIP1 are not (Figures 5 and S3). Interestingly, we also found less OPTN and TNIP1 recruitment to either LLOMe-or AMDE1-induced sites of LC3B lipidation compared to diABZI treatment, whereas p62, NBR1, and TAX1BP1 were recruited to LLOMe– and AMDE1-induced LC3B puncta/foci to more similar extents (Figures 5 and S3). NDP52 showed less recruitment in Monensin and LLOMe conditions, and similar levels of recruitment in AMDE1 conditions, compared to diABZI, but not as substantially as OPTN (Figures 5C and S3). Together these results show that autophagy receptors tend to be recruited to sites of LC3B-associated membranes induced by a variety of membrane damaging stimuli, but OPTN and TNIP1 recruitment appears to be more specific to STING activation.

**Figure 5.**
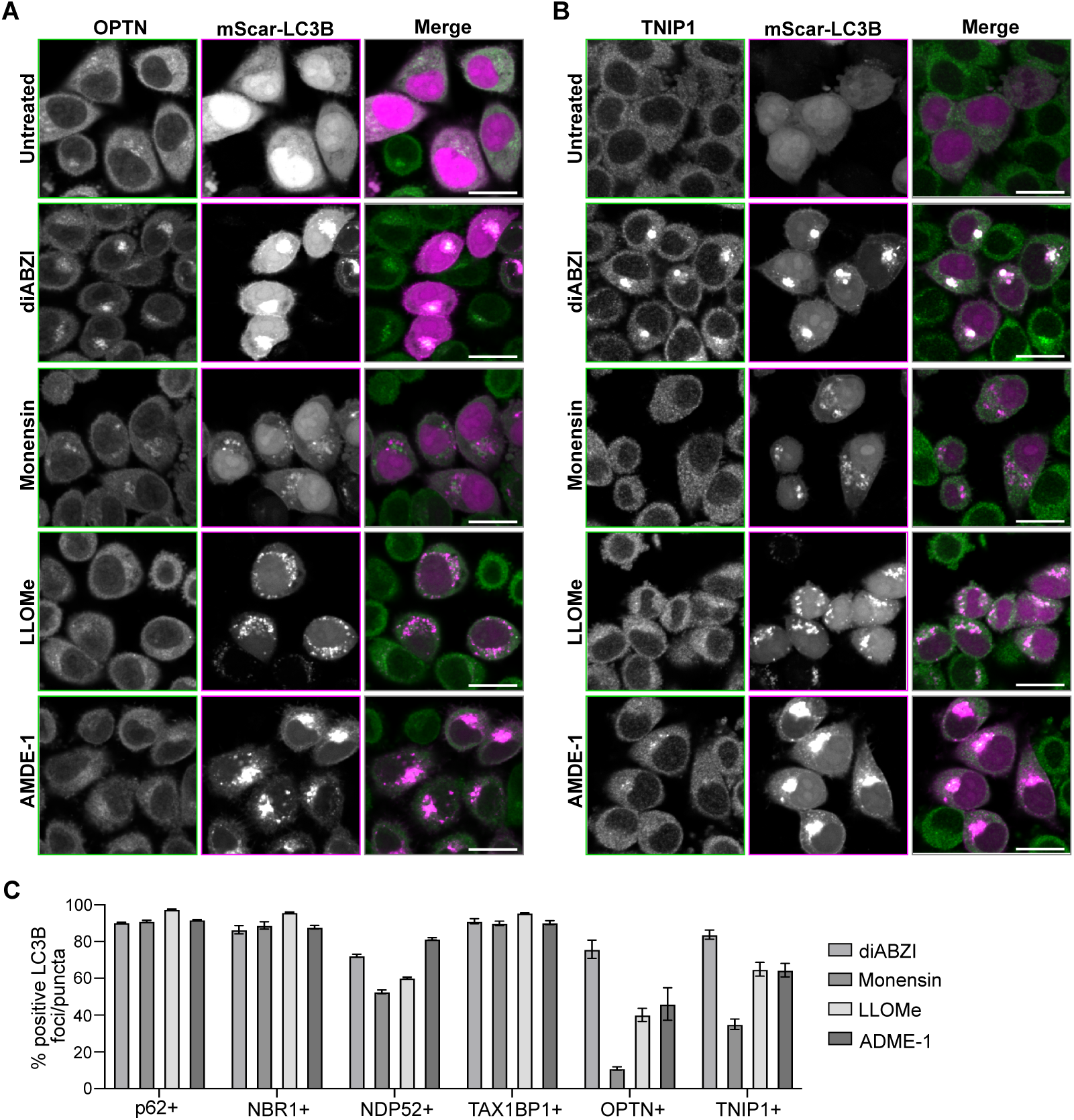
OPTN and TNIP1 are specifically recruited to LC3B associated membranes upon STING activation compared to other CASM-related stimuli. **A)** Representative spinning disk confocal images of WT HeLa^STING^ cells expressing mScarletI-LC3B (magenta) and immunostained for OPTN (green) after no treatment, diABZI (6 hr), Monensin (1 hr), LLOMe (1 hr), or AMDE-1 (6 hr). Scale bar indicates 20 µm. **B)** Representative spinning disk confocal images of WT HeLa^STING^ cells expressing mScarletI-LC3B (magenta) and immunostained for TNIP1 (green) after no treatment, diABZI (6 hr), Monensin (1 hr), LLOMe (1 hr), or AMDE-1 (6 hr). Scale bar indicates 20 µm. **C)** Quantification of images in panels A-E depicting fraction of LC3B puncta/foci positive for indicated proteins. Over 1000 cells were imaged for each condition.

While all five of these autophagy receptors contain LIR motifs, they vary in their ubiquitin binding domains – p62 and NBR1 have Ubiquitin Associated (UBA) domains, NDP52 and TAX1BP1 have ubiquitin-binding Zinc Finger (ZF) domains, and OPTN has a ZF immediately downstream of a UBAN (Ub-Binding domain in ABIN proteins and NEMO) domain (Johansen & Lamark, 2020). TNIP1 also contains a UBAN domain protein with LIR motifs that play a role in autophagy (Le Guerroué et al., 2023). Our data suggest that OPTN and TNIP1 are specifically recruited to LC3B foci that colocalize with STING in the context of STING activation relative to other damaging agents (Figures 5 and S3). To interrogate this recruitment without potential homodimerization with endogenous OPTN and TNIP1, we reconstituted OPTNKO and TNIP1KO cells, respectively, with corresponding mEGFP-tagged WT, LIR mutant, UBAN mutant, or LIR/UBAN double mutant proteins and assessed recruitment to LC3B foci following STING activation. WT and all mutant mEGFP-OPTN variants colocalized with LC3B at early timepoints following STING activation, however the UBAN mutant (D474N) and LIR/UBAN double OPTN mutants plateaued after approximately 2 hours, while the WT and the LIR mutant (F177A/E179A) continued to correlate more with LC3B at later times (Figures 6A-B and S4A). Assessment of mEGFP-TNIP1 WT and mutant variants, revealed substantially less co-localization of the TNIP1 UBAN (D472A) and double LIR/UBAN mutants with LC3B compared to the WT and LIR mutant (F83A/L86A and F125A/V128A) upon diABZI treatment (Figures 6C-D and S4B). These data indicate that recruitment of OPTN and TNIP1 to LC3B associated vesicles following STING activation is partially dependent and mostly dependent (respectively) on their UBAN domains and not their LIR motifs. The UBAN-independent OPTN recruitment observed may be explained by its TBK1-dependent recruitment (Figure 4, E-G), suggesting two potential mechanisms of OPTN recruitment to LC3B foci upon STING activation.

**Figure 6.**
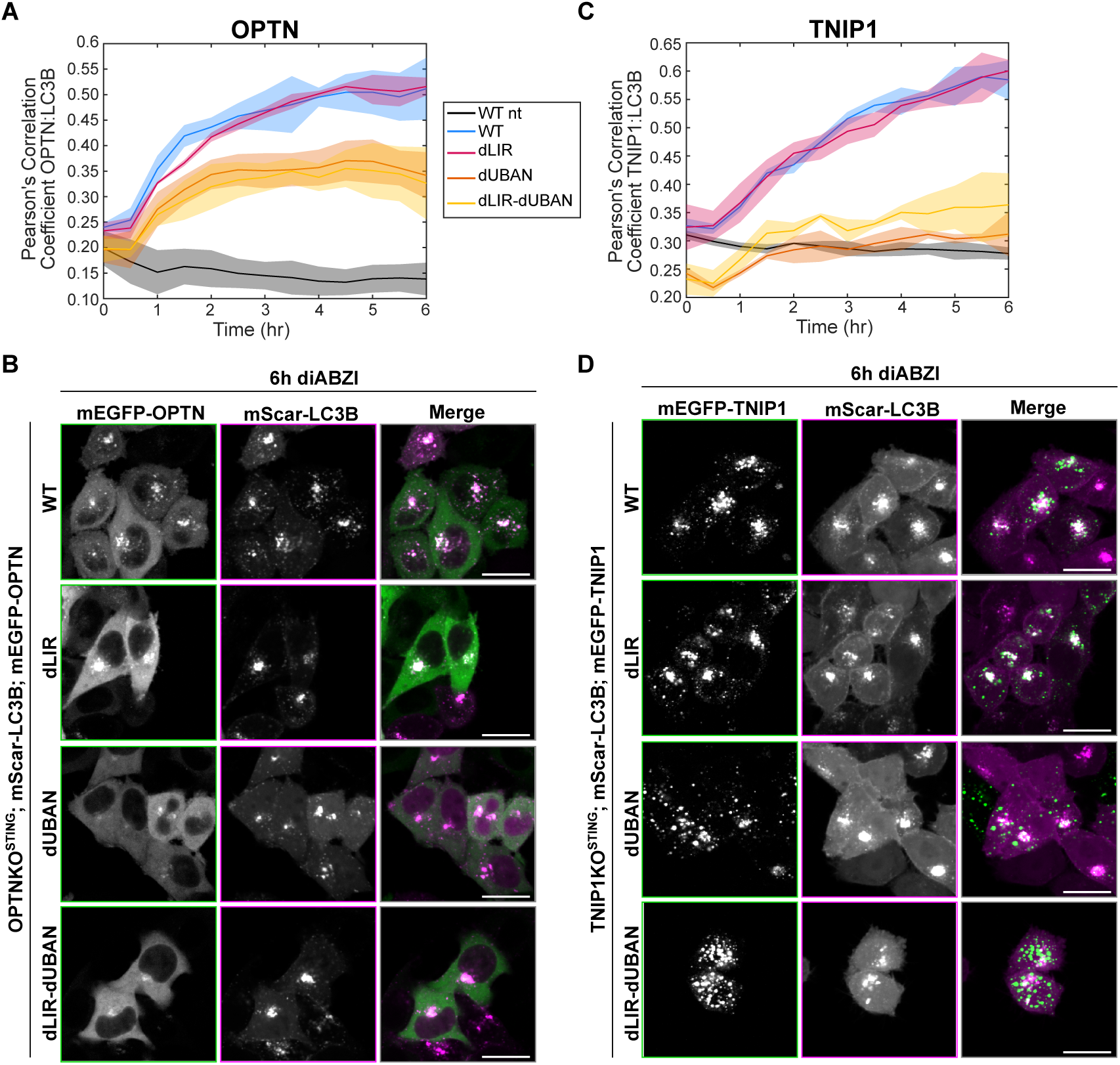
The UBAN domains, but not LIR motifs, are important for OPTN and TNIP1 recruitment to perinuclear LC3B foci upon STING activation. **A)** Pearson’s correlation coefficient of mScarletI-LC3B and mEGFP-OPTN-WT or mutants (dLIR(F83A, L86A, F125A, V128A), dUBAN(D472N), and dLIR-dUBAN (F83A, L86A, F125A, V128A and D472N)) plotted against time after treatment with diABZI or no treatment (nt). Shading represents standard deviation between wells. **B)** Representative spinning disk confocal images corresponding to panel A of WT HeLa^STING^ cells expressing mScarletI-LC3B (magenta) and mEGFP-OPTN (green) with indicated mutations after 6 hr diABZI. Scale bar indicates 20 µm. **C)** Pearson’s correlation coefficient between mScarletI-LC3B and TNIP1-WT or mutants (dLIR(F177A and E179A), dUBAN(D474N), and dLIR-dUBAN (F177A, E179A and D474N)) plotted against time after treatment with diABZI or no treatment (nt). Shading represents standard deviation between wells. **D)** Representative spinning disk confocal images corresponding to panel C of WT HeLa^STING^ cells expressing mScarlet-LC3B (magenta) and mEGFP-TNIP1 (green) with indicated mutations after 6 hr diABZI. Scale bar indicates 20 µm.

We have previously shown that STING activation leads to an increase in K63– and M1-linked ubiquitin chain formation, whereas Monensin treatment only induces K63-linked chain formation (Fischer et al., 2024). Because UBAN domains have been shown to bind M1 ubiquitin chains with a higher affinity than K63-linked ubiquitin chains *in vitro* (Herhaus et al., 2019; Nanda et al., 2011; Rahighi et al., 2009; Yoshikawa et al., 2009), we assessed TNIP1 and OPTN recruitment in HOIPKO cells, which lack M1 but not K63 chain synthesis (Fischer et al., 2024). Activation of STING by cGAMP in HOIPKO HeLa^STING^ cells showed similar recruitment of TNIP1 and OPTN compared to WT HeLa^STING^ cells (Figure S5, A-D), indicating that the UBAN dependent recruitment is not mediated by M1-linked/linear ubiquitin chains and may be more dependent on K63-Ub chains.

Both OPTN and TNIP1 have been reported to regulate interferon and NF-κB signaling (Cohen et al., 2009; Dziedzic et al., 2018; L. Gao et al., 2011; Gleason et al., 2011; Mauro et al., 2006; Medhavy et al., 2024; Meena et al., 2016; Munitic et al., 2013; Nagabhushana et al., 2011; Nakazawa et al., 2016; Nanda et al., 2011, 2016; O’Loughlin et al., 2020; Shinkawa et al., 2022; Su et al., 2019; Sudhakar et al., 2009; Zhou et al., 2011, 2023; Zhu et al., 2007). To assess if OPTN and TNIP1 may regulate immune signaling following STING activation, we knocked out OPTN and TNIP1 in the THP1 human monocyte cell line. Knockout of OPTN did not show significant effects on LC3B lipidation, the phosphorylation of STING, TBK1, or IRF3, IkBα degradation, nor the formation of M1-linked ubiquitin chains (Figure 7A), indicating OPTN is not required for activation of STING signaling. KO of OPTN similarly did not substantially affect STING-induced IRF3-or NF-κB-related gene expression (Figure 7, B-C), however the expression of the IFNβ stimulated gene, *IFIT3*, was significantly reduced (Figure 7B), potentially implicating a role for OPTN downstream of IRF3-induced IFNβ and consequential IFNβ receptor signaling induction of interferon stimulated genes. KO of TNIP1 in THP1 cells also did not present a defect in LC3B lipidation, however the phosphorylation of STING is increased in TNIP1KO cells at the 4-hour timepoint and there is less detectable IkBα compared to WT (Figure 7D), indicating IkBα may be more degraded and NF-κB activation enhanced. Strikingly, detection of both STING-induced IRF3-, interferon-, and NF-κB related gene expression in the TNIP1KO cells is significantly increased compared to WT (Figure 7, E-F). Thus, TNIP1 appears to negatively regulate both interferon– and NF-κB-mediated signaling downstream of STING.

**Figure 7.**
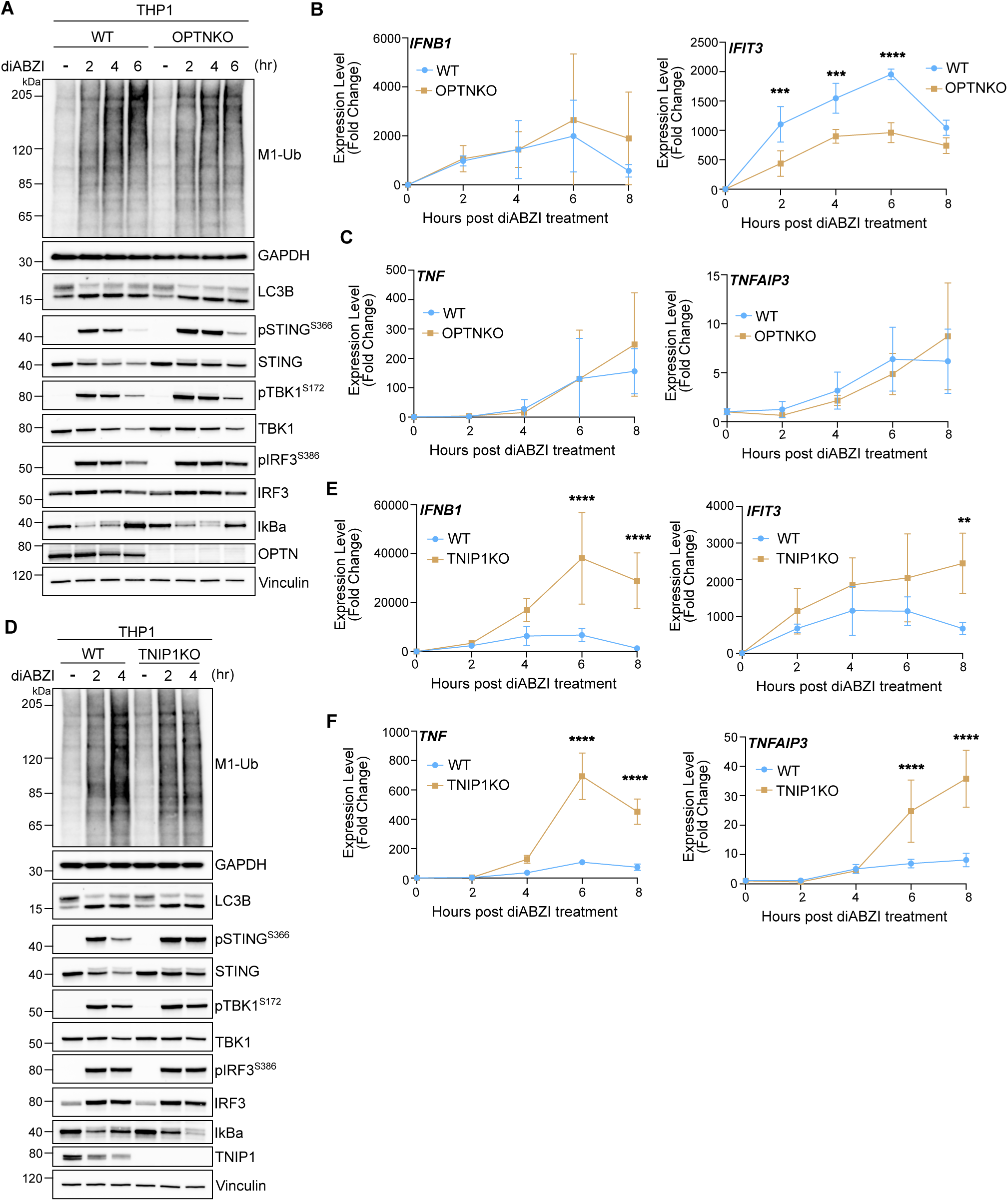
TNIP1 negatively regulates STING-mediated immune signaling. **A)** Representative immunoblots of indicated proteins detected in THP1 cell lysates from WT and OPTNKO cells prepared following treatment with 1 µM diABZI for 2, 4, and 6 hours. Immunoblotting was replicated in 3 independent experiments. **B-C)** Relative expression changes of indicated IRF3/interferon-related (B) and NFκB-(C) genes detected by quantitative RT-PCR in WT and OPTNKO THP1 cells treated with 1 µM diABZI for 2, 4, 6, and 8 hours. Quantification of relative expression is from 3 independent experiments analyzed at the same time. A 2-way ANOVA with a Sidak’s multiple comparisons test was performed on 2^-ΔΔCt^ values. Mean +/-s.d. n=3 *<0.05, **<0.01, ***<0.001, ****<0.0001. **D)** Representative immunoblots of indicated proteins detected in THP1 cell lysates from WT and TNIP1KO cells prepared following treatment with 1 µM diABZI for 2 and 4 hours. Immunoblotting was replicated in 3 independent experiments. **E-F)** Relative expression changes of indicated IRF3/interferon-related (E) and NFκB-(F) genes detected by quantitative RT-PCR in WT and TNIP1KO THP1 cells treated with 1 µM diABZI for 2, 4, 6, and 8 hours. Quantification of relative expression is from 3 independent experiments analyzed at the same time. A 2-way ANOVA with a Sidak’s multiple comparisons test was performed on 2^-ΔΔCt^ values. Mean +/-s.d. n=3 *<0.05, **<0.01, ***<0.001, ****<0.0001.

## Discussion

Activation of STING initiates an innate immune response after trafficking to the Golgi apparatus where it also induces ubiquitylation and LC3B lipidation at single membrane Golgi-derived vesicles (Fischer et al., 2020, 2024; Huang et al., 2025; Ishikawa et al., 2009; Saitoh et al., 2009). Here we find that the ubiquitin and LC3B binding proteins p62, NBR1, TAX1BP1, NDP52, Optineurin, and TNIP1 are recruited to LC3B and ubiquitin associated membranes following STING activation. Autophagy receptors are recruited to polyubiquitin chains conjugated to cargo during selective autophagy, linking the ubiquitinated cargo to autophagy machinery and facilitating their encapsulation by autophagosomes (Johansen & Lamark, 2020). p62 and NBR1, in particular, are important for the sequestration and clustering mechanism of ubiquitinated proteins (Sun et al., 2018; Turco et al., 2021; Zaffagnini et al., 2018) and organelles, like mitochondria (Narendra et al., 2010), peroxisomes, and endosomes (Deosaran et al., 2013; Mardakheh et al., 2010). Consistent with this known sequestering function, we found that p62 and NBR1 act redundantly to cluster LC3B-associated vesicles into perinuclear foci. We also found that LC3B clustering into foci is mediated by TBK1 kinase activity, although TBK1 is not required for the recruitment of p62/NBR1 to LC3B structures. These findings suggest that phosphorylation of p62 or NBR1 by TBK1 may regulate their sequestration activity, as Ser28 within the PB1 domain of p62 is a known phosphorylation site by TBK1 (Richter et al., 2016). Indeed, p62 has been shown to co-localize with LC3B upon STING activation, and its phosphorylation by TBK1 may play a role in STING degradation (Prabakaran et al., 2018). p62 and NBR1 have also been recently reported to play opposing roles in regulation of STING and interferon signaling in hepatic stellate cells and hepatocellular carcinoma phenotypes (Nishimura et al., 2024). Further study will be necessary to determine whether the clustering functions of LC3B– and Ub-associated Golgi-related membranes described here plays a role in the reported functions of p62/NBR1 in STING signaling.

Similarly to STING-mediated LC3B lipidation, LC3B can be conjugated to single membrane vesicles through the V-ATPase-ATG16L1 axis upon membrane damage, a process widely referred to as CASM (Figueras-Novoa et al., 2024; Fischer et al., 2020). We find that p62, NBR1, NDP52, and TAX1BP1 are also recruited to LC3B-associated structures upon treatment with CASM-related stimuli – the ionophore Monensin, the membranolytic polymer LLOMe, and the Golgi-damaging agent AMDE-1 (Durgan et al., 2021; Y. Gao et al., 2016; Thiele & Lipsky, 1990). Interestingly, both OPTN and TNIP1 had reduced recruitment to LC3B-associated structures in CASM conditions, distinguishing them from the other Ub– and LC3B-binding autophagy receptors in STING signaling. OPTN and TNIP1 are among the few proteins that bind M1-linked/linear polyubiquitin chains through their UBAN (Ub-Binding domain in ABIN proteins and NEMO) domains (Fennell et al., 2018). OPTN and TNIP1 also bind K63-linked polyubiquitin chains through the UBAN domain (Herhaus et al., 2019; Nanda et al., 2011). Mutations in the UBAN domain reduced both OPTN and TNIP1 recruitment to perinuclear LC3B foci following STING activation, revealing that these two proteins are recruited to the LC3B and ubiquitin foci downstream of STING activation in a ubiquitin-dependent and LIR-independent manner. TNIP1 appears to be recruited completely through its ubiquitin binding domain, whereas OPTN partially acts through the UBAN domain. We also found that TBK1 and its interaction with STING is important for OPTN recruitment. TBK1 has two coiled-coil domains (CCD) approximately 130 residues apart and binds to STING through CCD1 and OPTN through CCD2 (Morton et al., 2008; Runde et al., 2022; Zhang et al., 2019), potentially allowing for simultaneous interaction with both STING and OPTN at LC3B foci. Thus, OPTN can be recruited to LC3B– and Ub-associated foci by at least two modalities – the STING-TBK1 axis as well as its interaction with ubiquitin. Further we show that, while the UBAN domains are important for OPTN and TNIP1 recruitment to LC3B– and Ub-associated perinuclear foci, HOIP, the E3 ligase that synthesizes M1-linked ubiquitin chains upon STING activation (Fischer et al., 2024), is not required. These data indicate that binding to M1-linked/linear ubiquitin chains is dispensable for UBAN-dependent recruitment of OPTN and TNIP1 and may more so favor a model in which they bind K63-linked ubiquitin chains at LC3B foci following STING activation (Fischer et al., 2024).

TNIP1 and OPTN, and their binding to M1– and K63-linked polyubiquitin chains, are known to regulate both interferon and NF-κB related immune signaling through unclear mechanisms (Cohen et al., 2009; Dziedzic et al., 2018; L. Gao et al., 2011; Gleason et al., 2011; Mauro et al., 2006; Munitic et al., 2013; Nagabhushana et al., 2011; Nakazawa et al., 2016; Nanda et al., 2011, 2016; O’Loughlin et al., 2020; Shinkawa et al., 2022; Sudhakar et al., 2009; Zhu et al., 2007). Here we show that loss of OPTN had little impact on STING mediated NF-κB signaling. However, curiously, STING-mediated expression of *IFIT3*, an Interferon β-stimulated gene, was significantly reduced. These data indicate that the STING-TBK1 and/or UBAN mediated recruitment of OPTN is dispensable for upstream activation of STING and gene expression, however OPTN may have a function in signaling downstream of Interferon β. More interestingly, we discovered that TNIP1 may negatively regulate both Interferon and NF-κB – related immune signaling in the STING pathway. Gene expression mediated by both IRF3 and Interferon β, as well as NF-κB, were significantly increased with the loss of TNIP1 in THP1 monocytes upon STING activation. Increased IkBα degradation was detected at the 4-hour time point following STING activation, indicating that TNIP1 may limit NF-κB-related gene expression by negatively regulating the turnover of IkBα or a necessary step upstream. However, no clear changes in phosphorylation of IRF3 and TBK1 were detected by western blotting, suggesting that TNIP1 may be involved in a step in between TBK1 phosphorylation of IRF3 and IRF3-induced gene expression or in a feedback mechanism that shuts the signaling off. These results position TNIP1 as a negative regulator of STING-mediated interferon and NF-κB signaling similarly to its role in other immune pathways (Cohen et al., 2009; Dziedzic et al., 2018; L. Gao et al., 2011; Mauro et al., 2006; Medhavy et al., 2024; Nanda et al., 2011, 2016; Shinkawa et al., 2022; Su et al., 2019; Zhou et al., 2011, 2023). More work will be necessary to dissect the precise mechanisms in how OPTN and TNIP1 are playing their respective roles in innate immune signaling in the STING pathway.

## Material and Methods

### Cell Culture

HeLa and HEK293T cells (ATCC) were grown in DMEM supplemented with 10% FBS, 1x sodium pyruvate, and 1x GlutaMax at 37°C, 5% CO2, and 75% humidity. HeLa cells were passaged with 0.05% Trypsin–EDTA and aliquots frozen in BamBanker at –80°C. THP1 (Synthego-ATCC) cells were cultured in Roswell Park Memorial Institute (RPMI) 1640 (ATCC).

### Cell Lines

Knockout TBK1 and OPTN HeLa cells as well as p62/NBR1/NDP52/TAX1BP1/OPTN 5KO HeLa cells were generated previously in Lazarou et al. (Lazarou et al., 2015). p62, NBR1, p62/NBR1DKO HeLa cells were generated in Sarraf et al. (Sarraf et al., 2020). TNIP1KO HeLa cells were generated in Le Guerroue et al. (Le Guerroué et al., 2023). HOIPKO HeLa cells were generated in Fischer et al. (Fischer et al., 2024). OPTNKO THP1 cells were generated with lentivirus prepared as described below with the Lenti CRISPR v2 plasmid and the guide sequence GAUUUGAGGAGCUUUCGGCC. Following lentiviral transduction, THP1 cells were selected with 1 µg/mL puromycin for 2 days and then subcloned in 96 well plates. TNIP1KO THP1 cells were generated as a pooled knockout population by Synthego using the guide sequences: CGGCCAGCUCUUCACCCACC, GCAUCAGGUUGCCGUCCUCA, and CUGCUCAUUCUUG-UGCACCU, then subcloned in 96 well plates. Subcloned cells were assessed by Western blot for loss or truncation of the protein of interest.

### Cloning and Viral Transduction

All cloning was performed by PCR and Gibson assembly (New England Labs) in either pHAGE, pBMN, or pTRIP lentivectors and verified by Sanger sequencing or whole plasmid sequencing. Lentivectors were transfected into HEK293T cells with Tat1b, Rev1, MGMp2, and VSVg helper plasmids using polyethylenimine (PEI). After two days the viral inoculum was filtered (0.45µm) and added to cells of the appropriate genetic backgrounds for gene transduction as previously described (Wang, 2020).

### FACS Analysis and Sorting

To establish full coverage of stable protein expression, we sorted cells for a low STING expression by fluorescence signal of BFP (BFP-P2A-STING) and expression of mScarletI-LC3B. All cell lines were sorted using the same gates for comparable results. After sorting, cells were grown in Penicillin-Streptomycin-Neomycin (PSN) Antibiotic Mixture (GIBCO) containing media for five days. No experiments were performed in antibiotic-containing media.

### Treatment Conditions

Cells were treated with 1 µM diABZI (MedChem Express HY-112921B) for 6 hr unless otherwise stated. Cells were treated with 10 µM GSK8612 (MedChem Express HY-111941) simultaneously with diABZI when stated; washouts were performed with a single wash of DMEM before adding diABZI-containing media back to the well. Cells were treated with 120 µg/mL cGAMP (ChemieTek CT-CGMAP) for 8 hr, 10 µM AMDE-1 (MedChem Express HY-119137) for 6 hr, 100 µM Monensin (Sigma-Aldrich M5273) for 1 hr, or 1 mM LLOMe (Cayman Chemical Company 16008) for 1 hr.

### Live Cell Imaging

Cells were imaged on a Nikon Ti-2 CSU-W1 spinning disk confocal system operated by Nikon elements AR microscope imaging software using a 40x water objective (NA 1.15) with the automated TI2-N-WID water immersion dispenser. The microscope is equipped with a live cell chamber that was set to 37°C and 50% CO_2_ with constant humidity. Cells were imaged in P96-1.5H-N, Coverslip #1.5H 96-well imaging plates (Cellvis). mEGFP was excited with a 488 nm laser and assessed with a 525/36 nm emission filter; mScarletI was excited with a 561 nm laser and assessed with a 605/52 nm emission filter; and Hoescht was excited with 405 nm laser and assessed with a 455/50 nm emission filter.

### Immunofluorescence

Cells seeded in either 96-well imaging plates described above or 8-well chamber slides (Cellvis C8-1.5H-N) were fixed with 4% warm paraformaldehyde (PFA) in PBS for 10 minutes, washed three times with PBS, permeabilized with 0.5% Triton-X100 in PBS for 5 minutes, blocked with 3% goat serum, 1% BSA, and 0.1% Triton-X100 in PBS for one hour. Cells were then incubated overnight in 1:200 or 1:500 dilution of primary antibodies in blocking solution on a rocker at 4°C. Cells were washed three times, incubated with a 1:1000 dilution of secondary antibodies for one hour at room temperature in blocking solution on a rocker, then washed three times before imaging. Antibodies for immunofluorescence used were as follows: TNIP (Proteintech 15104-1-AP), OPTN (Proteintech 10837-I-AP), p62 (Abnova H00008878), NBR1 (Abnova 2-96 H00004077), NDP52 (Abcam ab68588), TAX1BP1 (Cell Signaling 5105S). Secondary antibodies labeled with Alexa 647, anti-mouse (Life Technologies A21235) or anti-rabbit (Life Technologies A21244), were used to visualize above proteins excited with a 640 nm laser and assessed with a 705/72 nm emission filter.

### Image Analysis

Image analysis was performed with custom MATLAB scripts. All images were background subtracted prior to segmentation and analysis. For Pearson’s Correlation Coefficient, cells were segmented from background using the intensity of mScarletI-LC3B and filtered by size, then segmented into single cells with a water shedding algorithm on a blurred image of either the mScarlet-LC3B channel or a merge of Hoescht and mScarletI-LC3B. Pearson’s Correlation was then performed on single cells, then the median value found between single cells for the entire image. A median value of each imaging site was found for the well and 3 wells were used for an average and standard deviation value.

To assess percent LC3B puncta or foci positive for signal in another channel, background subtraction and segmentation was performed as above, followed by a segmentation for punctate structures. A Gaussian blur was performed on the image to remove details smaller than the size of a cell, then the original (background subtracted) image was divided by the blurred image to generate an image with relatively equal intensities between cells. Bright pixels from the original (background subtracted) image and the equalized image were found with manual thresholds (held constant between images and conditions), then multiplied together to find common pixels in both. Segmented pixels were then multiplied by the cell mask before size exclusion was performed to remove noise and find structures of expected size – larger for foci (35 pixels) than for puncta (5 pixels). Each grouping of connected pixels representing an LC3B punctum or focus was then assessed in a loop to determine if positive for the other channel. A cellular region near the structure was found by morphological dilation and exclusion of puncta/foci and background pixels, to obtain a cellular region near the punctum or focus. The average intensity of the secondary channel (i.e. autophagy receptor) was then calculated for the punctum/ focus and the surrounding cellular region, and a ratio was found and assigned to the corresponding structure. Foci or puncta with a ratio greater than the manual threshold of 1.25 were considered positive for the secondary channel. All MATLAB scripts are available upon request. A schematic for this analysis is also included in Fischer et al. (Fischer et al., 2024).

To measure the dispersion of foci, images were background subtracted, cells segmented from background and underwent single cell segmentation as with the Pearson’s analysis above. In a loop for each cell with puncta or a focus/foci (determined by having a high enough number of non-zero elements in the puncta/focus mask), the weighted centroid of the LC3B intensity was found. Within each measured cell, another loop was run on all pixels of increasing distance from the centroid to measure LC3B intensity; pixels outside the cell segmentation were not counted. Pixels of the same distance around the centroid were averaged together for the cell. The LC3B values arranged by distance to the centroid for each cell were then normalized to the mean intensity for that cell. The median value for all cells in the well was found, then the average value and standard deviation between wells calculated and used to generate a plot. Each experiment was repeated at least three times with each condition in triplicate.

### Immunoblotting

THP1 cells were counted on an automated cell counter and plated at 0.8 x10^6^ cells/mL for each condition in 12-well or 6-well plates for experiments. Following indicated treatments, cells were pelleted by centrifugation at 400xg for 1 minute, washed with ice-cold HBSS and re-pelleted, and then frozen on dry ice. Cell pellets were stored at –80°C until prepared for western blotting. In preparation for immunoblotting, cell pellets were briefly thawed, lysed in ice-cold 1x LDS sample buffer (Thermofisher) supplemented with cOmplete Protease Inhibitor cocktail (Roche), and boiled immediately at 99 °C for 10 min. Protein quantitation per sample was obtained using Pierce BCA protein assay kit (Thermo Fisher). Dithiothreitol (DTT; Sigma) was added to each sample at a final concentration of 100 mM just before gel loading.

For general immunoblotting procedures, 20 µg of total protein per sample was loaded into wells of 4–12% Bis-Tris gels (Genscript) and separated in MES or MOPS buffer ran at 65 V for 30 min, then 125 V until the dye front reached the bottom of the gel. Separated proteins were then transferred onto 0.45 µm Nitrocellulose membranes in Bio-Rad Transblot Turbo Transfer Buffer with 20% EtOH using the Bio-Rad Trans-Blot Turbo Transfer system (semi-dry). Membranes were blocked in 5% milk dissolved in 0.05% Tween 20 in TBS (TBST) at room temperature (RT) for 1 h and thoroughly washed prior to primary antibody incubation in 3-5% BSA prepared in PBST at 4° C overnight. Membranes were then thoroughly washed in TBST and incubated in HRP conjugated secondary antibodies raised against the appropriate species in 5% milk TBST for 1 h at room temperature. Following final washes, HRP signal was developed using either Amersham ECL Prime (Cytiva) or SuperSignal West Femto ECL (Thermo Scientific) and detected using a ChemiDoc Imaging System (Bio-Rad). Images were analyzed using ImageLab (Bio-Rad).

A modified version of the Newton et al., 2012 protocol (Newton *et al*, 2012) was used for detection of M1-linked ubiquitin chains. 15-20 µg total protein was loaded into 3-8% Tris-Acetate gels and separated in tris acetate buffer (Invitrogen) at 100 V for 30 minutes and then 150 V until the dye front reached the bottom of the gel. Transfer of the separated proteins onto 0.45 µm nitrocellulose membranes was performed at 30 V for 2 hours in Tris-Glycine buffer (Towbin formulation) supplemented with 10% MeOH using the XCell SureLock blot module system (semi-wet). Membranes were blocked overnight at 4°C in 5% milk dissolved in TBST. Membranes were thoroughly washed in TBST and incubated in primary antibodies in 5% milk prepared in TBST for 1 hour and 15 minutes at room temperature. Membranes were then washed in TBST, incubated in HRP conjugated secondary antibodies raised against the appropriate species in 5% milk TBST for 1 hour at room temperature, and washed again in TBST. HRP signal was developed using SuperSignal West Femto ECL and detected and analyzed as described above.

### Quantitative Real-Time PCR

Quick-RNA MiniPrep Plus kit (Zymo Research) was used to extract total RNA from 5-10×10^5^ THP1 cells prior to reverse transcription with a High-Capacity cDNA Reverse Transcription Kit (Thermo Fisher Scientific). Equal amounts of cDNA from each biological sample, mixed with the corresponding primers at a final concentration of 0.5 µM, were prepared for qPCR using SYBR Green Master Mix (Thermo Fisher Scientific) and a CFX384 real-time system/C100 Touch Thermal Cycler (Bio-Rad). The Ct value of the gene of interest was normalized to the b-actin Ct to calculate ΔCt for each biological sample. Each ΔCt was then normalized to the average ΔCt of untreated samples to generate the ΔΔCt value. The 2^-ΔΔCt^ formula was used to calculate the relative gene expression figures (Livak & Schmittgen, 2001).

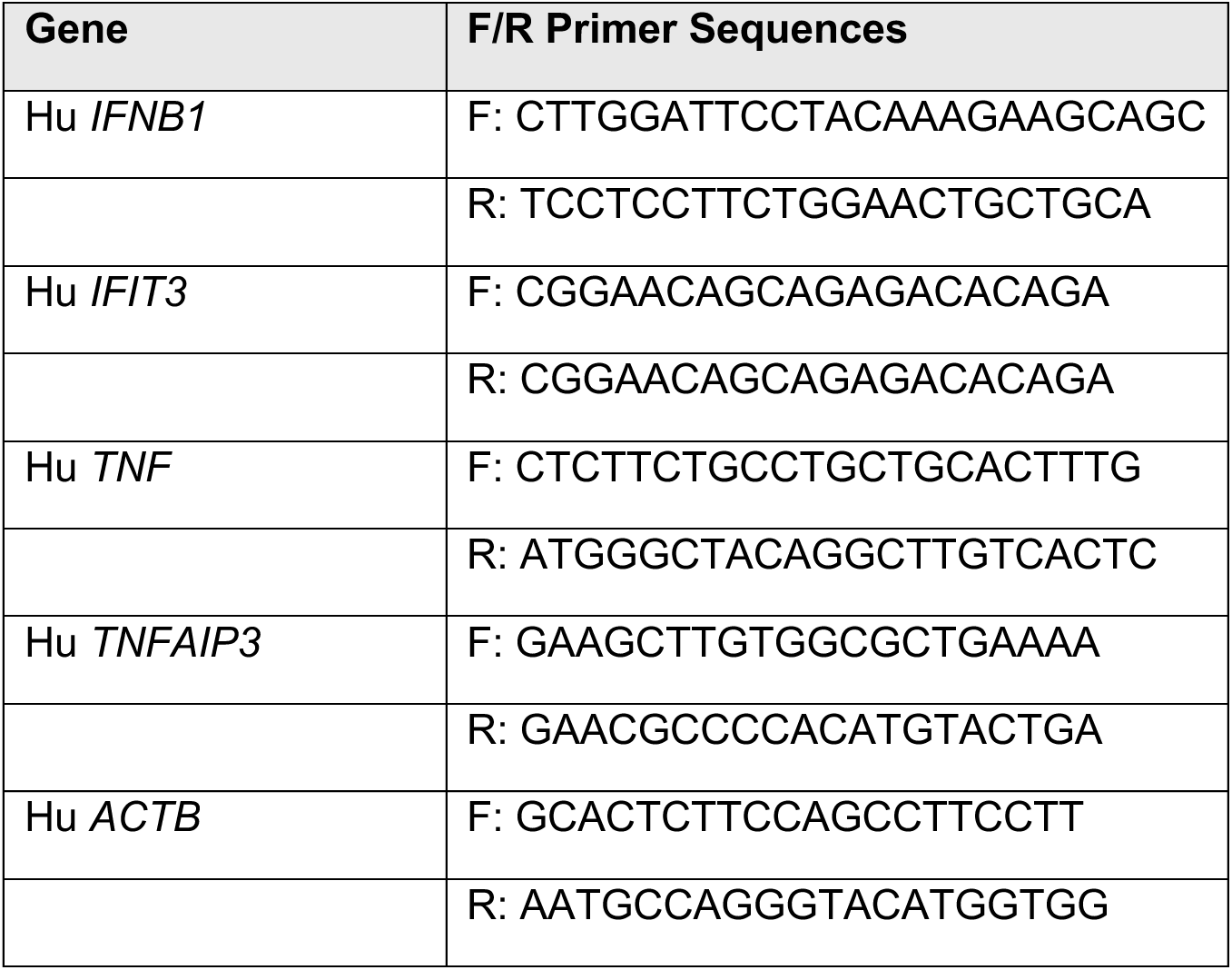

## Statistics

For imaging analysis, all p-values were generated with a two-way ANOVA using Tukey’s multiple comparisons. All error bars and filled shading represent standard deviation from the mean. For quantitative RT-PCR, 2-way ANOVA was performed on the 2^-ΔΔCt^ values with a Sidak’s multiple comparisons test. Statistical analyses were performed using Prism9 (GraphPad). Each experiment was performed at least three times unless otherwise stated.

## Acknowledgements

We would like to thank the NHLBI Flow Cytometry Core, the NINDS Light Microscopy Core, and the NINDS FACS core facility. This work utilized the computational resources of the NIH HPC Biowulf cluster. (http://hpc.nih.gov). Funding was provided by the Intramural Research Program (IRP) of the National Institute of Neurological Disorders and Stroke (NINDS)

## Figures Legends

**Figure S1.**
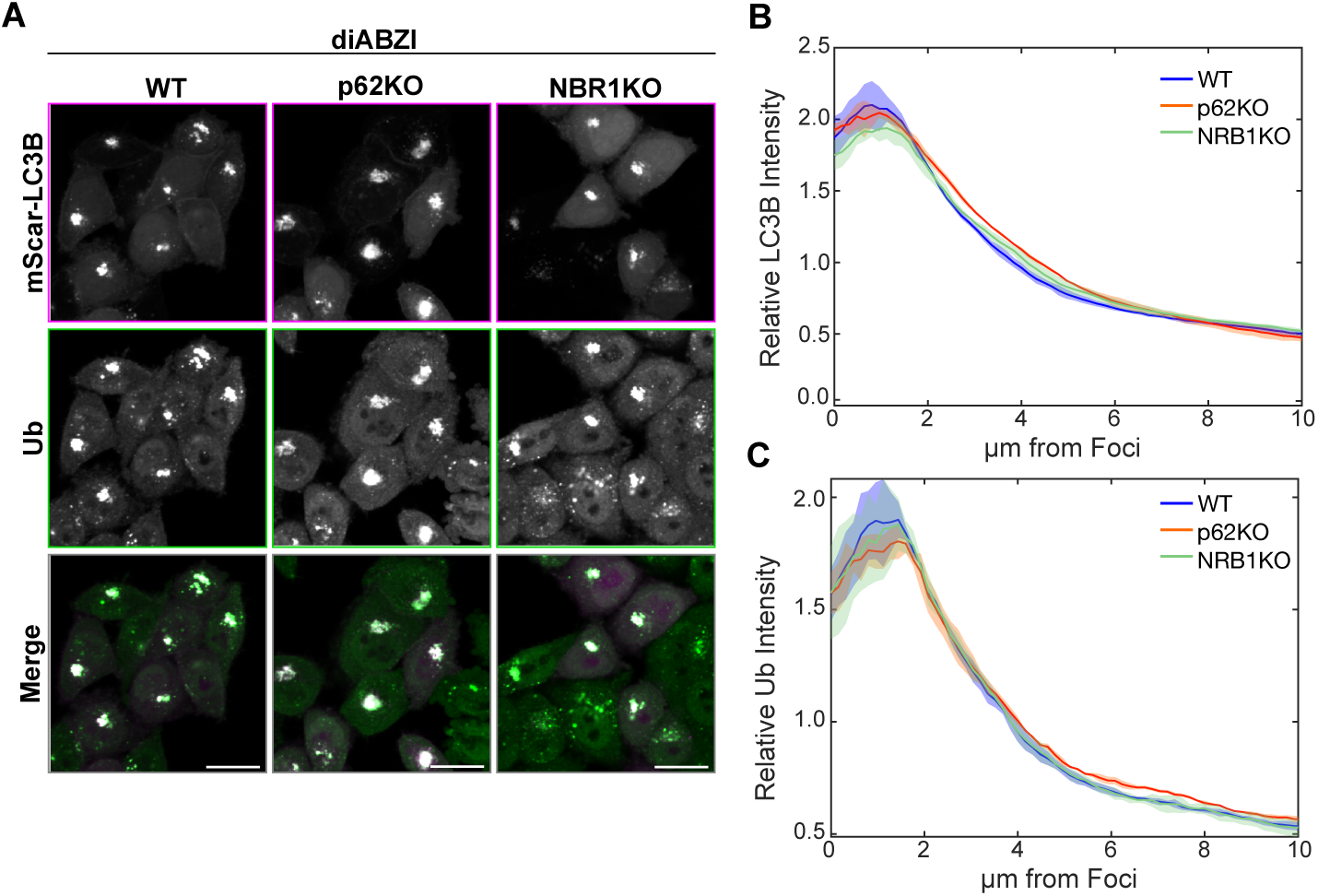
**A)** Representative spinning disk confocal images of WT, p62KO, or NBR1KO HeLa^STING^ cells expressing mScarlet-LC3B (magenta) treated for 6 hr with diABZI then fixed and immunostained for ubiquitin (green). Scale bars indicate 20 µm. **B-C)** Normalized intensity of LC3B (B) or signal within each cell plotted against distance to centroid of LC3B signal from cells displayed in panel A. Over 1000 cells were imaged for each condition.

**Figure S2.**
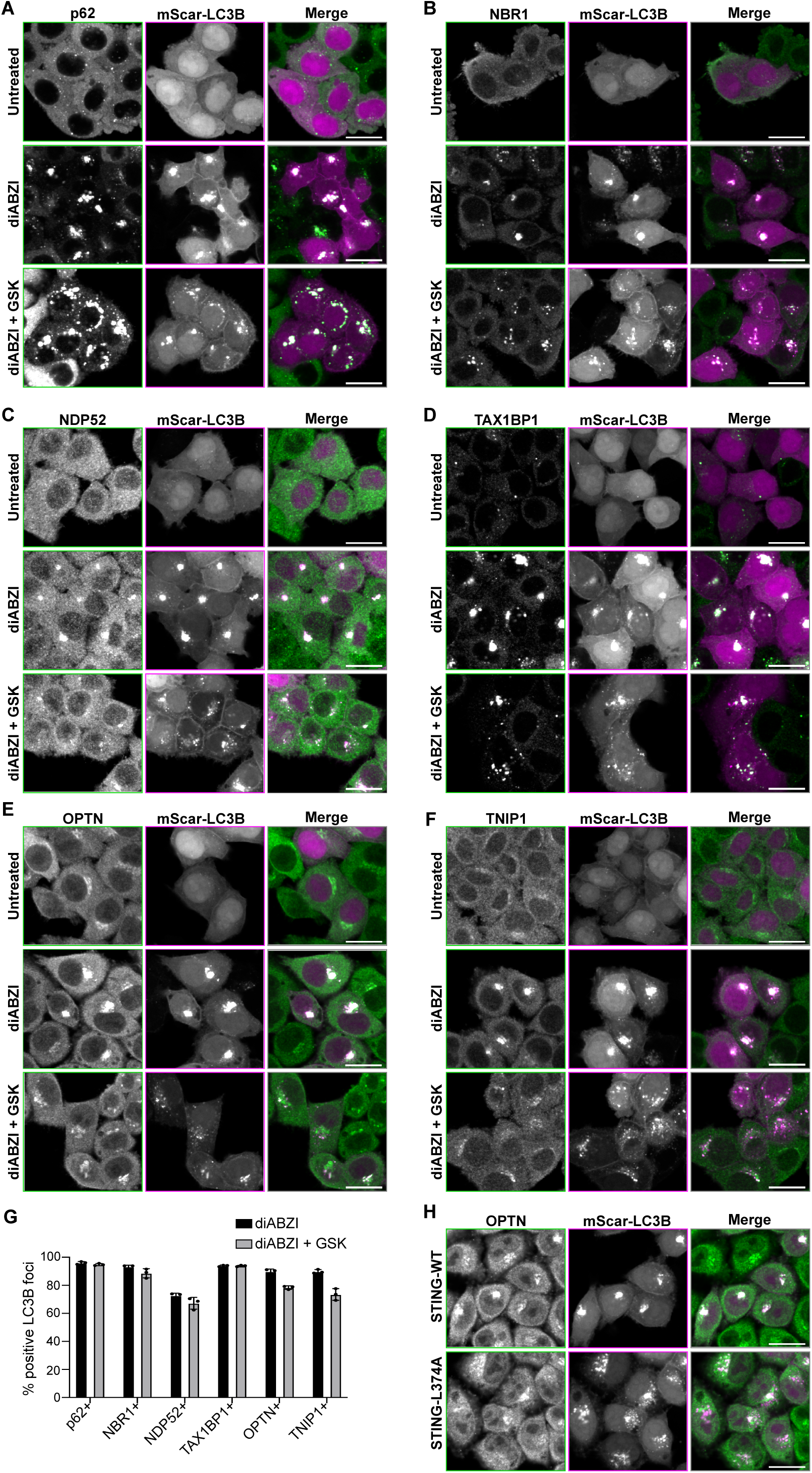
**A-F)** Representative spinning disk confocal images of WT HeLa^STING^ cells expressing mScarlet-LC3B (magenta) treated for 6 hr with diABZI or diABZI and TBK1 kinase inhibitor GSK, then fixed and immunostained for indicated proteins (green). Scale bars indicate 20 µm. G) Quantification of images in panels A-E depicting the fraction of LC3B foci positive for indicated protein. Over 1000 cells were imaged for each condition. H) Representative spinning disk confocal images of WT HeLa^STING^ or HeLa with STING_L374A_ expressing mScarletI-LC3B (magenta) treated for 6 hr with diABZI, then fixed and immunostained for OPTN (green). Scale bars indicate 20 µm.

**Figure S3.**
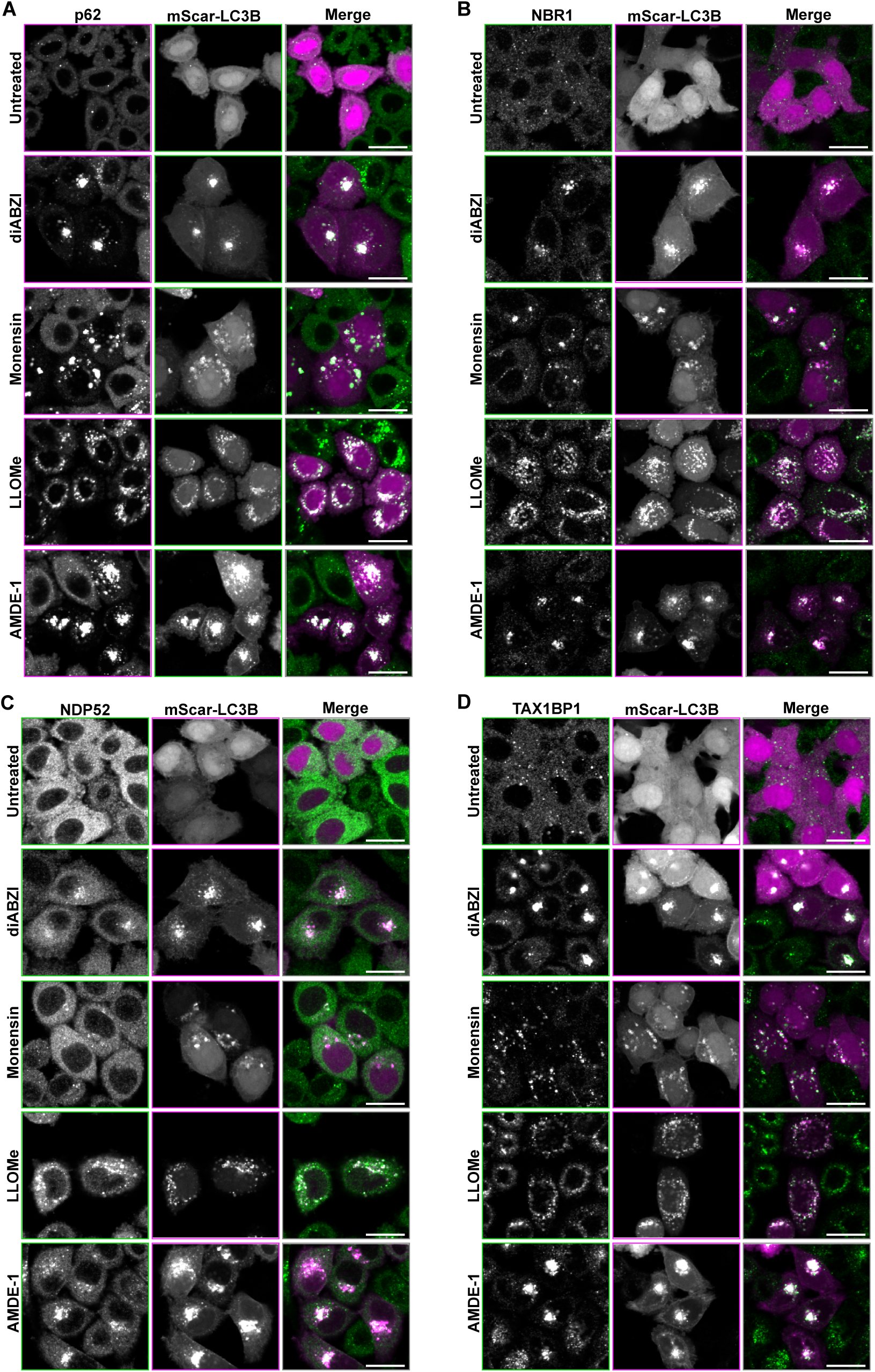
**A-D)** Representative images of WT HeLa^STING^ cells expressing mScarletI-LC3B (magenta) and immunostained for indicated proteins (green) after no treatment, diABZI (6 hr), Monensin (1 hr), LLOMe (1 hr), or AMDE-1 (6 hr). Scale bar indicates 20 µm.

**Figure S4.**
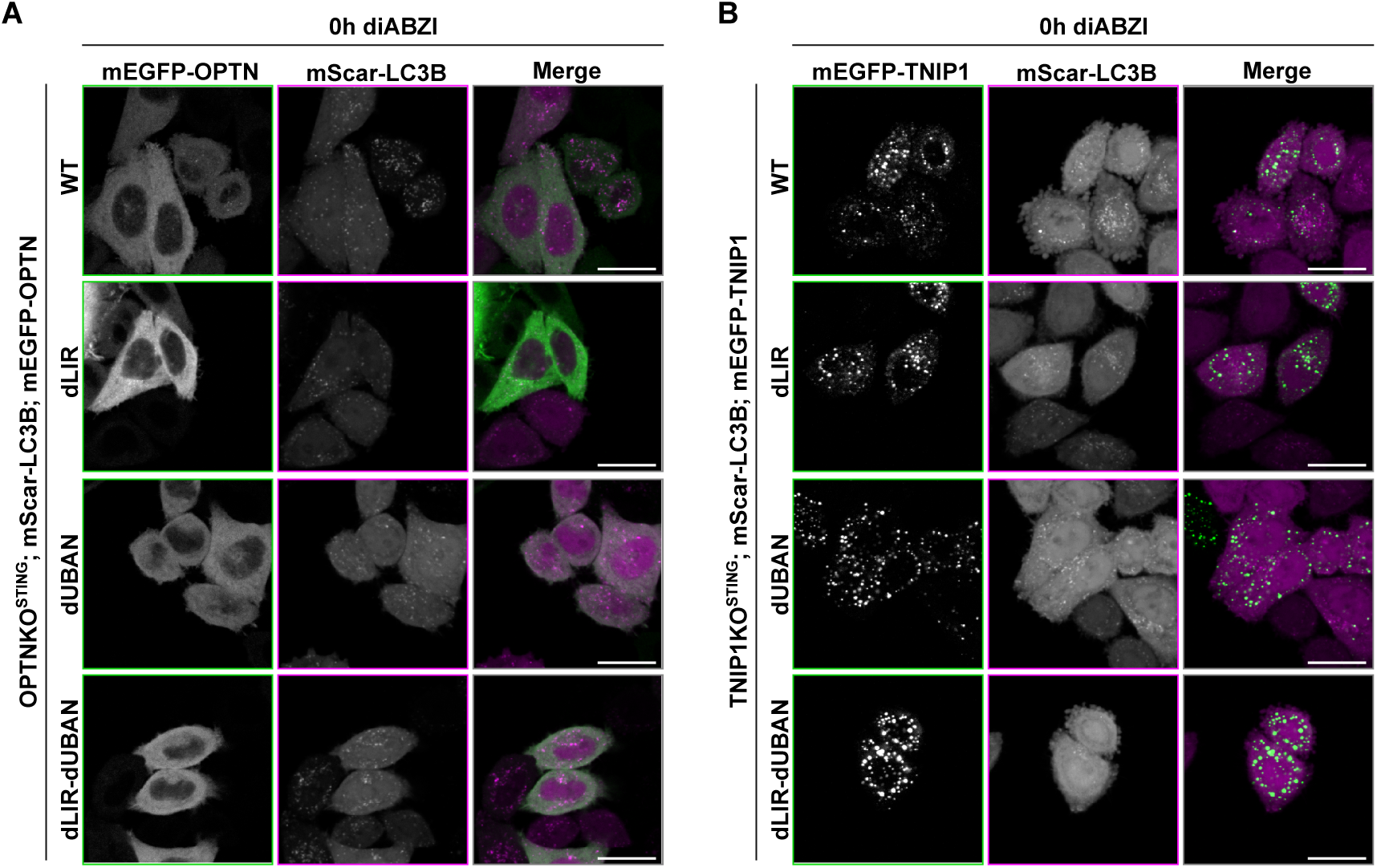
**A-B)** Representative spinning disk confocal images of OPTNKO (A) or TNIP1KO (B) HeLa^STING^ cells stably expressing mScarletI-LC3B and either mEGFP-OPTN-(**A**) or TNIP1-(**B**) WT, dLIR, dUBAN, or dLIR-dUBAN at the 0-hour and 6-hour time points represented in Figure 5, C-D following treatment with 1uM diABZI. Scale bar = 20 µm.

**Figure S5.**
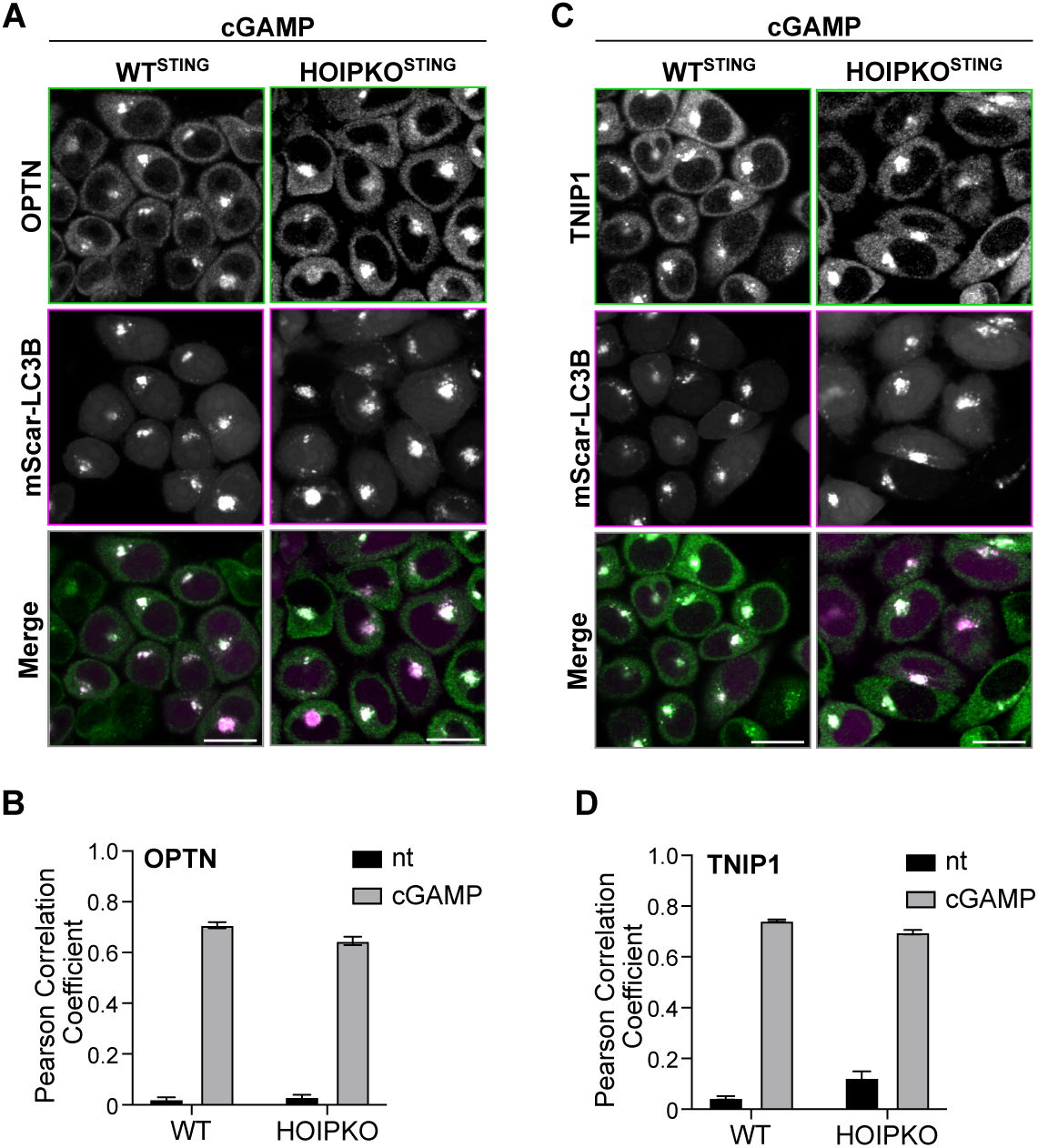
**A)** Representative spinning disk confocal images of WT and HOIPKO HeLa^STING^ cells stably expressing mScarI-LC3B (magenta) cells treated with 120 µg/mL of cGAMP for 8 hours prior to PFA-fixation and immunostaining for OPTN (magenta). Scale bar = 20 µm. **B)** Pearson correlation coefficient of mScarI-LC3B and endogenous OPTN from experiments represented in (C). Quantification is from 3 wells analyzed in the same experiment. Imaging was replicated in 2 independent experiments. **C)** Representative spinning disk confocal images of WT and HOIPKO HeLa^STING^ cells stably expressing mScarI-LC3B (magenta) cells treated with 120 µg/mL of cGAMP for 8 hours prior to PFA-fixation and immunostaining for TNIP1 (magenta). Scale bar = 20 µm. **D)** Pearson correlation coefficient of mScarI-LC3B and endogenous TNIP1 from experiments represented in (E). Quantification is from 3 wells analyzed in the same experiment. Imaging was replicated in 2 independent experiments.

